# Contig annotation tool CAT robustly classifies assembled metagenomic contigs and long sequences

**DOI:** 10.1101/072868

**Authors:** Diego D. Cambuy, Felipe H. Coutinho, Bas E. Dutilh

## Abstract

In modern-day metagenomics, there is an increasing need for robust taxonomic annotation of long DNA sequences from unknown micro-organisms. Long metagenomic sequences may be derived from assembly of short-read metagenomes, or from long-read single molecule sequencing. Here we introduce CAT, a pipeline for robust taxonomic classification of long DNA sequences. We show that CAT correctly classifies contigs at different taxonomic levels, even in simulated metagenomic datasets that are very distantly related from the sequences in the database. CAT is implemented in Python and the required scripts can be freely downloaded from Github.

## Introduction

Metagenomics involves direct sequencing of DNA from microbial communities in natural, clinical, or biotechnological systems. Limitations including the short length of DNA sequencing reads and the diversity of microbial communities made read mapping the ideal approach to answer the classical questions in metagenomics (Handelsman, 2004) i.e. "Who is there?" and "What are they doing?". Read mapping considers each read as an independent observation, whose taxonomic origin and functional class can be estimated by identifying the closest match in a reference database, and tallying these annotations across the full metagenomic dataset results in a taxonomic or functional profile of the microbial community (Wilke et al., 2016; Huson et al., 2016). Although this approach provides a high-resolution overview of the taxa and/or functions in a sample, the growing data volumes increasingly require heuristic sequence alignment tools. Together with the possibility that short reads may be ambiguously mapped to the reference sequences in the database, this may not always result in the most sensitive annotation. As a result, a persistent caveat of read mapping approaches is an abundance of "unknowns": sequences that do not yield a significant match in the reference database and are not reported (Dutilh, 2014).

In recent years, it has become increasingly clear that reference-based annotation of short-read metagenomes may miss an important fraction of the microbial and viral biodiversity in the environment (Dutilh et al., 2014; Brown et al., 2015). With increases in DNA sequencing volumes, metagenomics has moved from read mapping to sequence assembly, the additional coverage making it possible to assemble high-quality contiguous sequences from the metagenome. Moreover, with the advent of single molecule DNA sequencing platforms (third generation sequencing) such as Single Molecule Real Time (SMRT) sequencing (Eid et al., 2009) and Nanopore (Schneider & Dekker, 2012), long read lengths are becoming an attractive opportunity to complement short-read based metagenomes, albeit at a lower ecological resolution due to the smaller number of individual reads obtained with these platforms.

Longer sequences allow a less ambiguous annotation of the unknown queries, while the lower number of sequences to be annotated allow more sensitive, but computationally more demanding annota tion tools to be used. Typically, a single metagenome may consist of millions to hundreds of millions of 100-250 base pair sequences when using second-generation sequencing, versus just thousands to tens of thousands of 1,000-100,000 base pair assembled contigs or long-read sequences derived from metagenome assembly or from third-generation sequencing. These long sequences are often annotated with a best BLAST hit approach. While this approach tends to work well if the strains present in the microbial community have sequenced representatives in the database, we show here that this approach quickly breaks down as the metagenome contains sequences from more unknown organisms, i.e. organisms that are distantly related to the ones in the reference database.

Here we present the Contig Annotation Tool (CAT), designed to provide robust taxonomic classification of long sequences from unknown organisms, such as those found in a metagenome. We test CAT using simulated long metagenomic sequences with decreasing genomic similarity to the sequences in the reference database, showing that CAT outperforms the commonly used best BLAST hit approach. CAT is implemented in Python and the source code is freely available on Github.

## Methods

The Contig Annotation Tool (CAT) is a pipeline for robust and accurate taxonomic classification of long metagenomic sequences. The CAT pipeline is implemented in Python, and it uses third party programs for several steps. The source code and installation instructions are available at: https://github.com/DiegoCambuy/CAT.

Figure 1 displays the whole pipeline. CAT takes a set of DNA sequences as input, and first identifies open reading frames (ORFs) with Prodigal using default parameters (Hyatt et al., 2010). Second, protein translations of the predicted ORFs are queried against the NCBI non-redundant (nr) protein database (NCBI Resource Coordinators, 2016) to identify the top hits. Here, CAT takes into account all hits with a bitscore ≥90% of the highest bitscore for that ORF (all the parameter values are default values that can be adjusted by the user). The nr database is constantly updated, so we recommend using the latest version. Homology searches are performed with DIAMOND using default parameters (Buchfink, Xie & Huson, 2014). In order to derive a conservative taxonomic classification of an individual ORF, the taxonomic annotations of the top hits are compared, and their last common ancestor (LCA) determined according to NCBI Taxonomy. For each ORF, CAT records the taxon of their LCA (T) as well as the average of their bitscore values (B).

**Figure 1.**
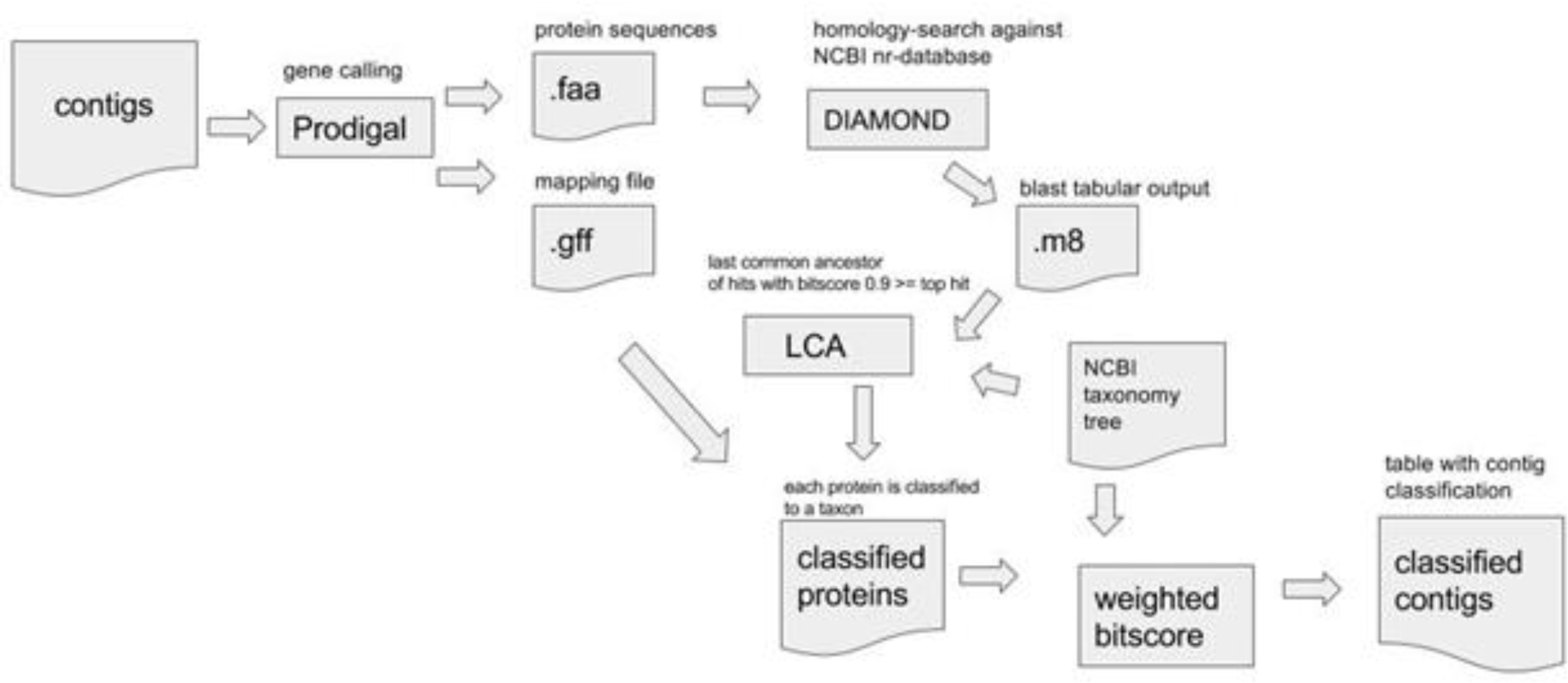
Outline of the CAT pipeline. For details see text.

By adjusting the cutoff value for how close the bitscore of the considered hits should be to the top hit (default: 90%), the user can tune the conservativeness of in the taxonomic classification of individual ORFs. As a result of this approach, ORFs that are highly conserved in taxonomically diverse species will receive a less specific taxonomic classification, i.e. they will be classified at a higher taxonomic levels (e.g. phylum, class, order). In contrast, ORFs that are very particular to one clade will receive a more specific classification, i.e. they will be classified at a lower taxonomic levels (species, genus, family). Finally, in order to address non-specific taxonomic annotations that are often present in the database, CAT has the default behavior to ignore hits if their taxonomic annotations are a parent of the taxonomic annotation of other hits of the same ORF.

Third, the individual taxonomic classifications are integrated across all ORFs to derive a robust classification of the full DNA sequence. To do this, the B values of all ORFs assigned to the same taxon are summed for each taxon (ΣBtaxon), where the taxonomic annotations of T and all of its parents in the taxonomic lineage are considered. For example, if the top hits of a given ORF share a LCA T at the taxonomic level of family, then their average bitscore value B would also be considered in the ΣBtaxon score of all ancestor nodes of T at the levels of order, class, and phylum. At the same time, CAT also calculates the maximum achievable ΣB value, ΣBmax, by summing the B values of all ORFs on the entire sequence. Finally, CAT assesses the ΣBtaxon values of all the identified taxa, and assigns the contig to that taxon if the value of ΣBtaxon is ≥0.5 times ΣBmax. At this default bitscore cutoff factor, a given sequence can be assigned to at most one taxon at each level (phylum, class, order, family, genus, and species). Howe ver, if the cutoff is lowered below 0.5 by one of the user options, it is theoretically possible that more than one taxon reaches the bitscore threshold. In this case, the query sequence is assigned to the taxon with the higher ΣBtaxon value.

### Testing data

We tested the performance of CAT by using a testing dataset with DNA sequences of varying length, whose taxonomic annotation was known. To create this benchmarking dataset, we randomly selected from the Genbank database(NCBI Resource Coordinators, 2016) 689 complete bacterial genomes, each from a distinct genus. These genome sequences were fragmented into 13,311 non-overlapping subsequences ranging from 175 to 399,980 base pairs in length. Prodigal identified 2,308,934 ORFs on these sequences.

Since the genomes used to create the benchmarking dataset were derived from the database, a straightforward CAT annotation according to the pipeline outlined above will easily identify the correct taxonomic annotation (see Results section). This is equivalent to a situation where a strain that is found in the metagenome has been sequenced, and its annotated sequence is present in the reference database. However, CAT was designed to also classify metagenomic sequences that are only distantly related to those in the database. Thus, to test the performance of CAT with increasingly unknown sequences by using our *in silico* dataset, we devised an approach where the top DIAMOND hits were increasingly ignored after the database search step. Specifically, hits were ignored if the sequence identity to the query in the aligned region exceeded a cutoff of 98% (well-studied metagenome), 78% (intermediate), or 58% (highly unknown sequences).

### Best BLAST hit

To determine the annotation of sequences in the benchmarking dataset by using the best BLAST hit approach, sequences were queried against the nr database with blastn (Altschul et al., 1990). Sequences were assigned to the taxonomic annotation of the best non-self hit. Self hits were removed based on the sequence identifier, i.e. not based on sequence identity as above, so that closely related organisms could be retained among the blastn hits.

## Results

To illustrate how CAT classifies a given DNA sequence, we describe one fictional but illustrative example in detail. In the example shown in Figure 2, the DIAMOND homology searches of the protein translations of five ORFs identified on the DNA sequence yielded four, zero, three, four, and four hits, respectively. For the first ORF, the top DIAMOND hit is present in *Escherichia coli* and the hit had a bitscore of 239. The same homology search yielded two other hits with bitscores ≥90% of that value: *Vibrio cholerae*(bitscore: 224) and *Klebsiella pneumoniae* (bitscore: 218). The bitscore of the fourth hit in Bacteroides fragilis was below the 90% threshold and this hit is ignored. Thus, CAT assigns to the first ORF an average bitscore value B = 227, and lists T = *Gammaproteobacteria* as its taxonomic classification, i.e. the LCA of *E. coli*, *V. cholerae*, and *K. pneumoniae*. The second ORF has no hits, and therefore does not contribute to the classfication of the sequence. For the third ORF, the LCA of the two hits T is reported as *Nitrospira marina*, because *Nitrospirae* is within the parental lineage of *N. marina*, and is thus ignored. This default behavior of CAT is implemented to address inspecific database annotations, and it can be optionally turned off. The fourth and fifth ORFs are assigned to the LCA of the taxa identified among the protein hits within 90% of their highest bitscore, similarly to the first ORF.

**Figure 2.**
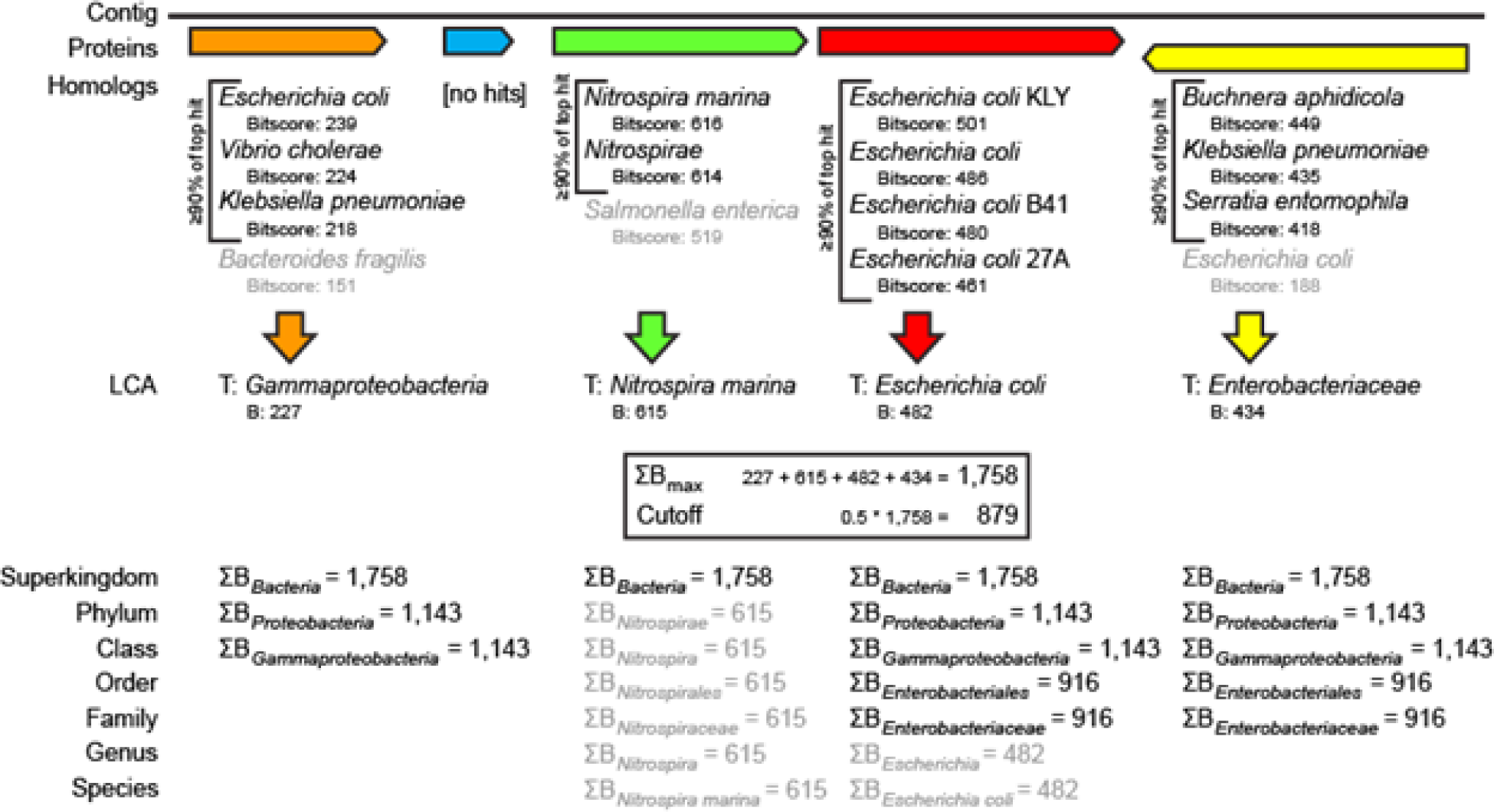
Illustrative fictional example of sequence classification by CAT. For details see text.

Next, the cutoff for reliable assignment of the contig to a given taxon is set at 0.5 times ΣB_max_ where ΣB_max_ is the sum of the B values of all ORFs and 0.5 is the default bitscore cutoff factor. In the example in Figure 2, ΣB_max_=1,758 and the cutoff is 879. CAT then assesses ΣB_taxon_ for taxa at all taxonomic levels, and in the example above the contig is assigned to *Bacteria* (ΣB_*Bacteria*_=1,758), *Proteobacteria* (ΣB_*Proteobacteria*_=1,143), *Gammaproteobacteria* (ΣB_*Gammaproteobacteria*_=1,143), *Enterobacteriales* (ΣB_*Enterobacteriales*_=916), and *Enterobacteriaceae* (ΣB_*Enterobacteriaceae*_=916), but not to *Nitrospirae* (ΣB_*Nitrospirae*_=615) or any of the levels below, or to *Escherichia* (ΣB*Escherichia*=482) or *E. coli*. Note that in the example in Figure 2, a best BLAST hit approach might have selected *Nitrospira marina* as the annotation for the contig, while the CAT pipeline provides a more robust annotation.

### Performance

Next, we assessed the performance of CAT on a benchmarking dataset of more than 13 thousand long DNA sequences with a known taxonomic origin. These sequences varied widely in length, and were sampled from complete genomes from 689 distinct genera. We benchmarked the performance by recording the accuracy and specificity of the classifications at each taxonomic level.

First, we tested the taxonomic annotation of the sequences by using the best BLAST hit approach, that is commonly used to classify unknown sequences. Interestingly, even after conservatively removing self hits, still about 80% of the sequences were wrongly classified by this approach at the species and genus levels (Figure 3, left bars). This not only reflects the sparse sampling of many genera, but also the fact that local alignment can result in a small region of the query being aligned (Figure 4).

**Figure 3.**
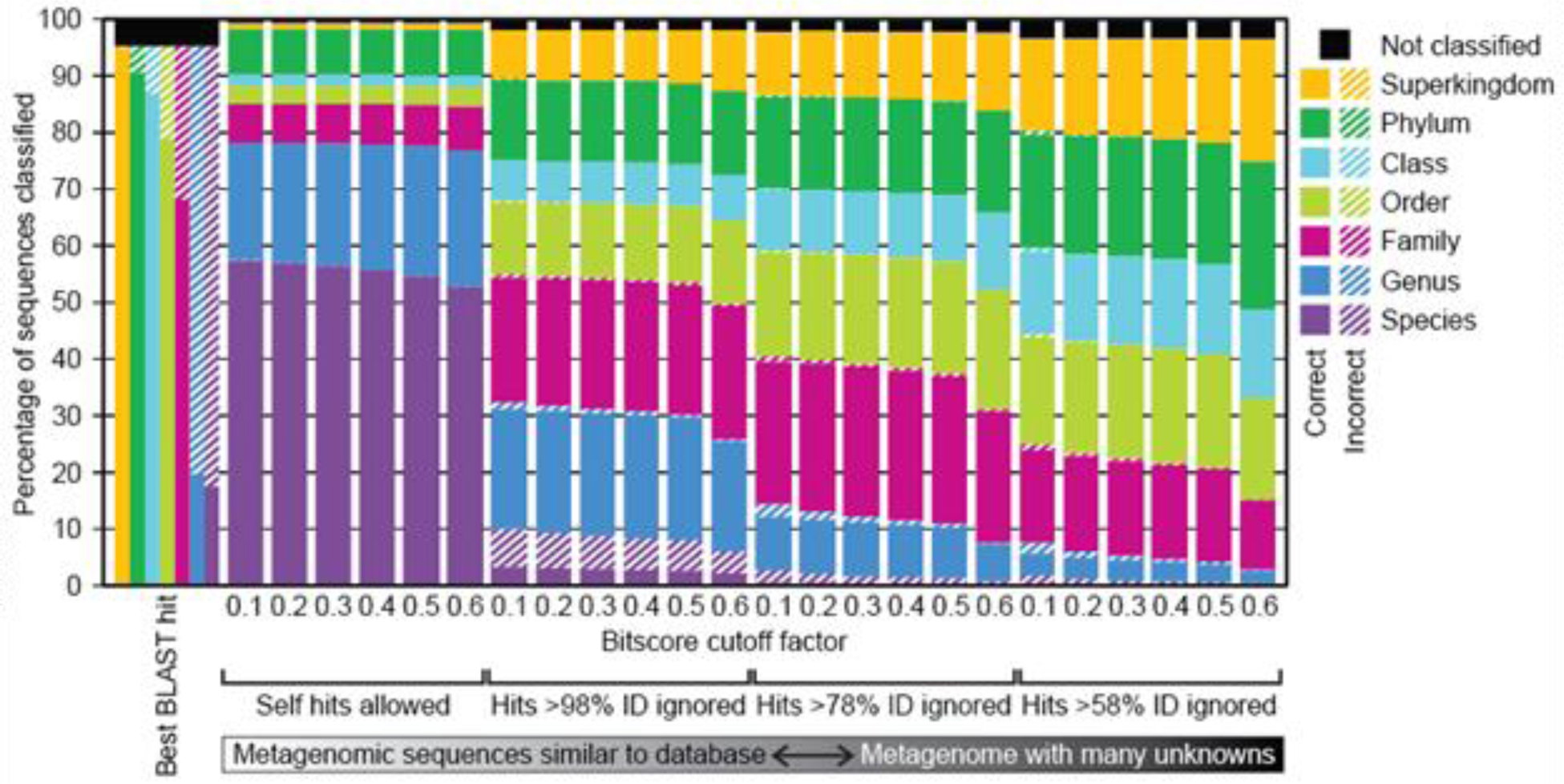
Classification of 13,311 sequences in the benchmarking dataset by using the best BLAST hit approach (left bars, self hits excluded) and CAT with six different bitscore cutoff factors (remaining stacked bars). The results are organized in four sets of six bars, representing increasingly unknown sequences (see Methods), where self hits were allowed, or removed at sequence identity (ID) >98% (well-known sequences), >78% (intermediate), or >58% (highly unknown sequences). For each taxonomic level, incorrectly assigned sequences are indicated as a red bar above that level. Unclassified sequences, i.e. those where no ΣB_taxon_ was higher than the bitscore cutoff, are indicated in black.

**Figure 4.**
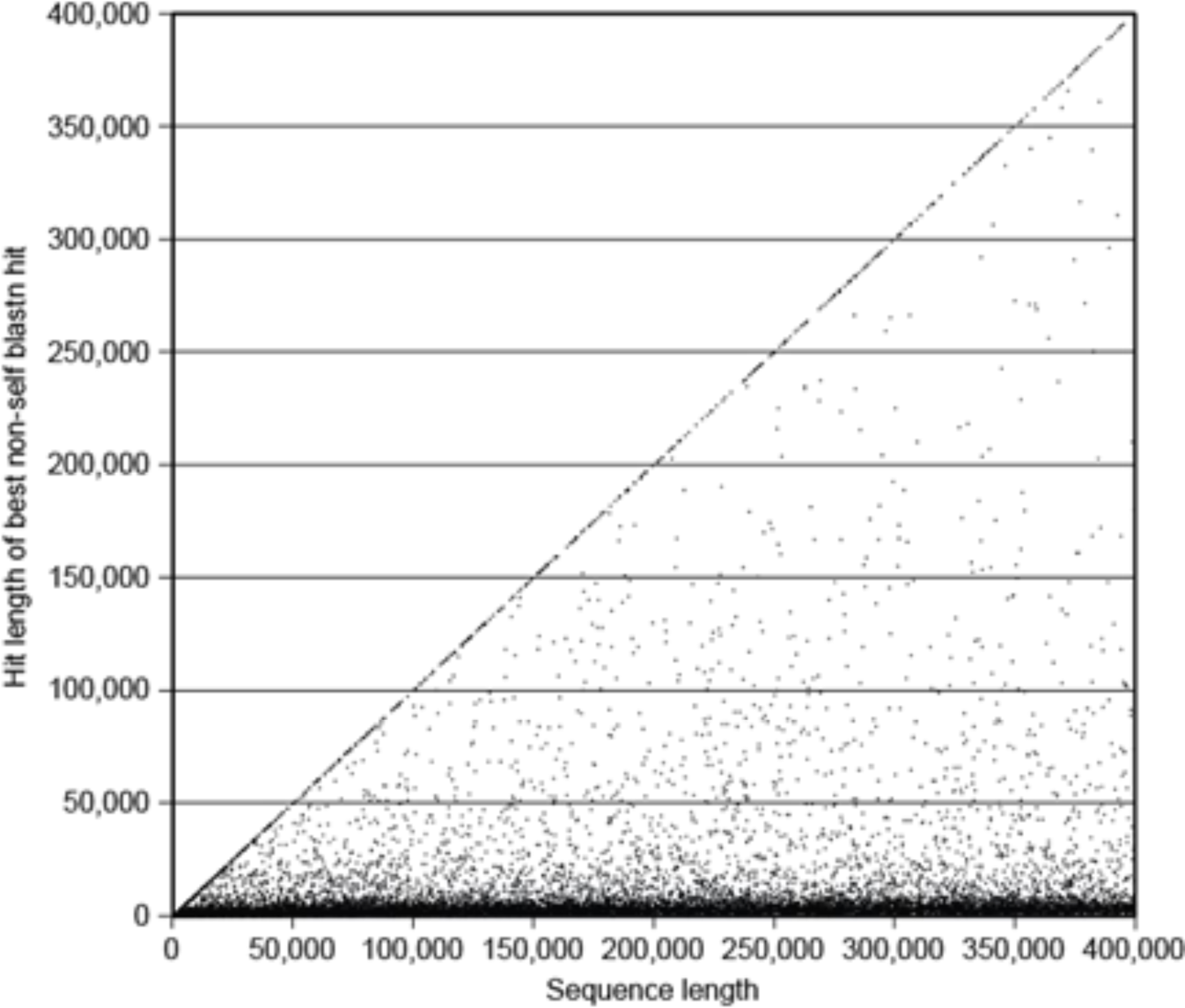
Length of the best non-self blastn hit for 13,311 DNA sequences in the benchmarking dataset. The accuracy of taxonomic classifications according to this approach is shown in Figure 3 (left bars).

In contrast to the best BLAST hit approach outlined above, CAT does not classify sequences based on a single best hit. Rather, CAT first determines the most likely taxonomic annotation of every individual ORF found on the sequence, and then assigns the full sequence to a taxon if this taxon is identified in enough of the ORFs, leading to a robust classification for the whole sequence. As expected, there were virtually no incorrectly classified sequences if self hits were not removed from the DIAMOND output (Figure 3), reflecting the case where the strain that is present in the microbial community has been sequenced, and is present in the database. The bitscore cutoff factor did not have a large influence on the level of taxonomic annotation. More than 40% of the sequences could not be annotated to the species level, indicating that for ORFs on these sequences, a substantial portion of the DIAMOND hits consisted of close homologs (here, ≥90% identity) in other taxonomic groups. As we discuss below, this conservative approach leads to a highly robust classification of metagenomic sequences derived from less well-studied environments, i.e. where the query is more distantly related to the sequences in the database.

Next, we filtered the most similar sequences from the DIAMOND hits, ignoring hits with >98%, >78%, and >58% identity in the aligned region. This reflects cases where the strains that are present in the microbial community are increasingly distantly related to those present in the database. As a result, CAT classifies the query sequences at increasingly lower taxonomic resolution. Since highly similar sequences from, e.g. the same species are no longer present among the hits, the contig can only be assigned to the next taxonomic level, e.g. genus or higher. Importantly, because excluding highly similar DIAMOND hits results in classification at a higher taxonomic level, the overall fraction of correctly annotated sequences increases from 93.1% to 98.3% (at bitscore cutoff factor 0.5). This is consistent with previous reports that higher taxonomic levels can be annotated more accurately (Randle-Boggis et al., 2016). The fraction of sequences that were not classified, i.e. when no ΣBtaxon reached the reliable assignment cutoff, increased from 1.7% to 3.2% with increasingly unknown query sequences (bitscore cutoff factor 0.5).

## Conclusions

We presented CAT, a bioinformatic pipeline to taxonomically classify long DNA sequences. CAT exploits homology information of all encoded proteins, leading to a robust classification for a long query sequence. We showed that thanks to this feature, CAT outperforms the frequently used best BLAST hit approach, that often misclassifies a sequence due to short, spuriously high scoring hits. Although annotations become less taxonomically resolved, we argue that this is realistically the case in many microbial communities, where the microbes that are found in a metagenome are only distantly related to the sequences in the database, and can thus only be reliably assigned to higher level taxa. When longer query sequences are available, such as contigs in the case of metagenome assemblies, or long sequencing reads from recent single-molecule sequencing platforms, CAT is a robust solution for taxonomic annotation.

